# TorchLIMIX: GPU-accelerated multivariate genome-wide association studies

**DOI:** 10.64898/2026.04.29.721342

**Authors:** Bibiana M. Horn, Zoran Nikoloski, Christoph Lippert

## Abstract

**Summary:** We introduce TorchLIMIX, a GPU-accelerated PyTorch implementation of the LIMIX multivariate genome-wide association study pipeline. By leveraging batched GPU linear algebra, TorchLIMIX achieves speedups of up to two orders of magnitude over the original CPU-based implementation while maintaining numerically equivalent results and full concordance of significantly associated loci. In simulation studies, replacing the default initialization of the genetic covariance factor with a QR-based strategy reduces genomic inflation factors to near-unity values under the common and interaction effect null hypotheses, ensuring well-calibrated type I error control. Applying TorchLIMIX to metabolic traits of *Arabidopsis thaliana* measured in two experiments uncovered 37 additional associated SNPs at the same significance threshold used in the original univariate GWAS.

**Availability:** The TorchLIMIX pipeline is openly available on GitHub at https://github.com/bi-horn/torchLIMIX. An adapted version of the multivariate association testing part of the original LIMIX pipeline is available at https://github.com/bi-horn/LIMIX_modified.

**Supplementary information:** Supplementary materials are included with this submission.

## Introduction

Genotype-by-environment (G*×*E) interactions capture the phenomenon whereby genotypes differ in their phenotypic response to environmental variation, reflecting underlying differences in phenotypic plasticity (2; 16). Here, ‘phenotype’ refers to the same trait measured across different environments, enabling the study of phenotypic plasticity. To understand phenotypic plasticity, it is essential to jointly analyze multiple (correlated) phenotypes using multivariate genome-wide association studies (GWAS) and to accurately model GxE effects—a central focus of quantitative genetics (6; 16). Linear mixed models (LMMs) provide a robust framework for multivariate GWAS (4; 10; 18). By jointly modeling genetic and residual covariance across phenotypes, LMMs account for the correlation structure among traits, which increases statistical power relative to separate univariate tests (14; 18). Population stratification is controlled by including a kinship matrix that captures genome-wide relatedness among individuals (17; 9). Within this framework, structured hypothesis tests can distinguish shared effects that act uniformly across phenotypes from phenotype-specific and G*×*E effects that vary between environments (11; 4). LIMIX introduced an efficient framework for multivariate linear mixed models by exploiting Kronecker structure in the covariance matrix (11). However, as datasets and phenotype panels grow, the cost of covariance learning and genome-wide association testing becomes a practical bottleneck. Graphics processing units (GPUs), while best known for their role in deep learning, offer benefits that extend beyond model training. Frameworks such as PyTorch (12) enable GPU-accelerated tensor operations and highly parallel execution, which speed up both covariance parameter estimation and per-SNP association testing. Based on this, we introduce TorchLIMIX, a GPU-accelerated implementation of the LIMIX multivariate testing framework (11) in PyTorch. We assess its scalability compared to the original NumPy-based pipeline, referred to here as NumpyLIMIX, and show that improved initialization of the genetic covariance matrix enhances both calibration and statistical power, and that multivariate analysis of correlated phenotypes uncovers associated loci not identified by univariate GWAS.

## Methods and Materials

### Multivariate linear mixed model

TorchLIMIX models phenotypes *Y ∈* ℝ^*n×p*^ for *n* individuals across *p* phenotypes using a multivariate LMM (11):

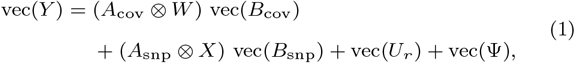

where *W ∈* ℝ^*n×c*^ contains *c* covariates, *X ∈* ℝ^*n×*1^ is the standardized dosage vector for the candidate SNP, *A*_cov_ and *A*_snp_ are trait-design matrices, and *B*_cov_ and *B*_snp_ the corresponding effect-size matrices. The genetic random effects are distributed as *U*_*r*_ ~ ℳ 𝒩 (**0**, *K*_stable_, *C*_0_), where *K*_stable_ denotes the variance-normalized genetic relatedness matrix and *C*_0_ ∈ ℝ^*p×p*^ the genetic covariance between phenotypes. Residual variation is captured by Ψ ~ℳ 𝒩 (**0**, *I*_*n*_, *C*_1_), with *C*_1_ ∈ ℝ^*p×p*^ the corresponding error covariance. For the genetic covariance factor *L*_0_ (where *C*_0_ = *L*_0_*L*^*⊤*^_0_), we evaluate two initialization strategies: a default method that sets all entries of *B*_0_ to one, and a QR-based approach that yields a well-conditioned initial covariance with unit eigenvalues. Full details on parameterization, optimization, and numerical stabilization are provided in Supplementary Information, Section 1 and Figure 1a. The framework supports five hypothesis tests: the *common effect* test for variants with uniform effects across all phenotypes, the *any effect* test for variants affecting at least one phenotype, and the *specific effect* test for single-phenotype associations. The *specific vs. common* and *any vs. common* tests assess deviations from a shared effect, thereby capturing interaction effects. Testing for these multivariate association patterns is achieved by appropriate definition of the SNP effect design matrix *A*_snp_ under both null and alternative hypotheses (see Supplementary Table S1).

**Figure 1.**
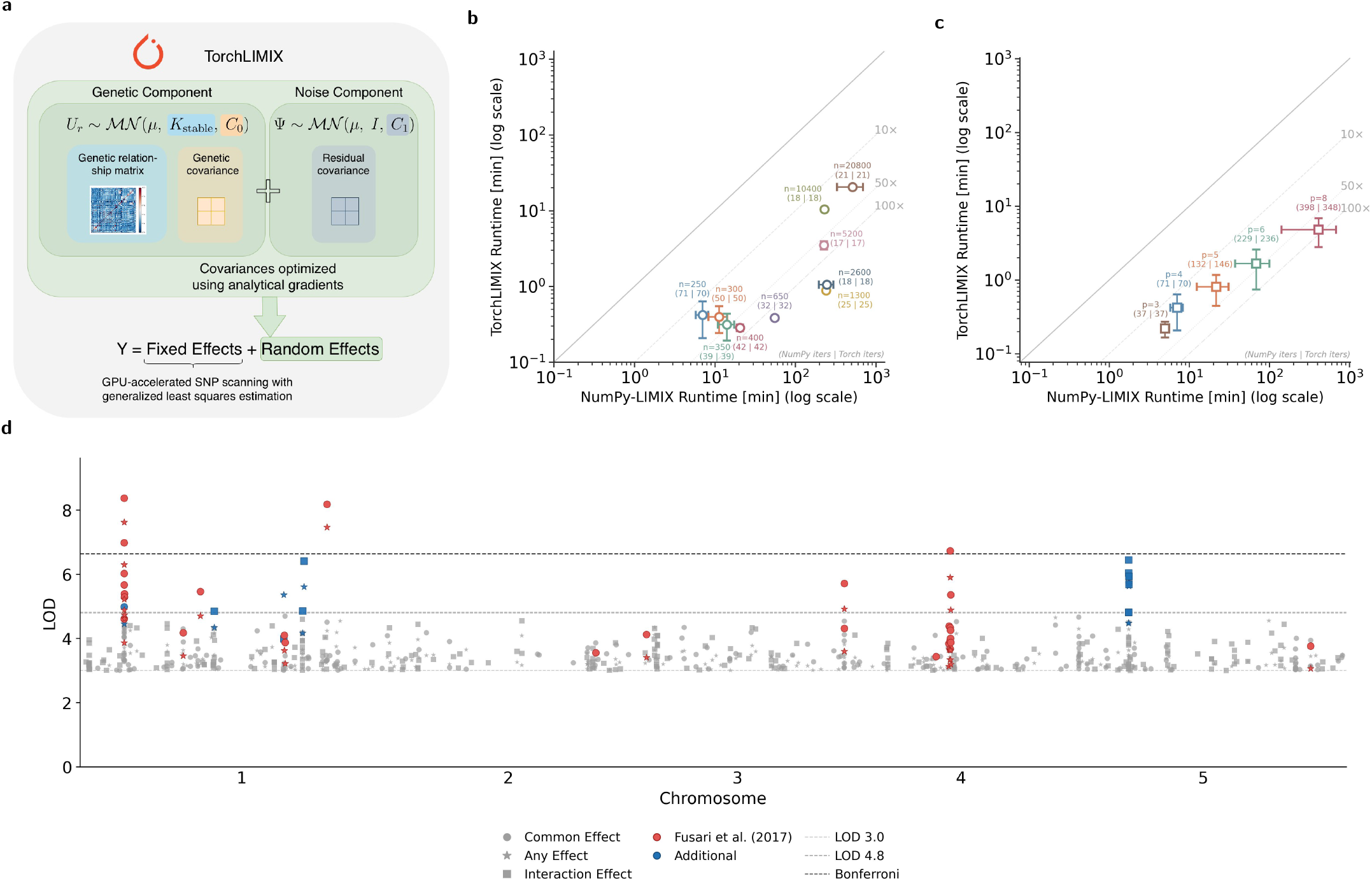
TorchLIMIX achieves scalable GPU based multivariate GWAS, recovers known and identifies additional associated loci. (a) The TorchLIMIX pipeline estimates genetic and residual covariance matrices using analytically derived gradients. The genetic random effects *U*_*r*_ follow a matrix normal distribution ℳ 𝒩 (0, *K*_stable_, *C*_0_), where *K*_stable_ is the variance normalized genetic relatedness matrix and *C*_0_ is the genetic trait covariance. The error covariance is modeled as *C*_1_ ⊗ *I*, capturing correlated error across traits and independent errors across samples. These covariance structures are incorporated as random effects into a LMM, and SNP effect sizes are estimated via GPU parallelized generalized least squares. (b) Scatter plot comparing NumPyLIMIX and TorchLIMIX runtimes across sample sizes (*n*) with four phenotypes. Each point represents the mean runtime over 10 timed runs (error bars show *±*1 sample standard deviation); distance below the diagonal indicates the speedup achieved by TorchLIMIX. Annotations show the median iteration count for each implementation. (c) Same comparison across phenotype counts (*p*) with 250 samples. NumPyLIMIX results for *p ≥* 10 are omitted as they exceeded the 100-hour wall-clock budget. (d) Manhattan plot across all five chromosomes of common (circles), any effect (stars), and any vs. common (squares) test statistics for the phenotype aINV. Red markers indicate loci previously identified by Fusari et al.; blue markers indicate novel associated loci. Horizontal lines mark LOD thresholds of 3, 4.8, and 6.63 (Bonferroni).

### *In silico* study with simulated phenotypes

Genotype data for *Arabidopsis thaliana* accessions were obtained from the HapMap diversity panel (8) in PED/MAP format and converted to PLINK binary format using PLINK v1.9 (13; 5). Minor allele frequency filtering of 10% was applied. From this dataset, we sampled *n* individuals with genotype matrix **G** ∈ ℝ^*n×s*^ (*s* SNPs) and simulated multivariate phenotypes **Y** ∈ ℝ^*n×p*^ across *p* environments following the generative model of Casale et al. (3) (Supplementary Information, Section 2). The proportional change in effect sizes across environments is controlled by a parameter, denoted by *η* (Supplementary Equations 2 and 3; Supplementary Table S2)). The polygenic background, hidden confounding, and noise components were sampled from matrix-normal distributions (Supplementary Table S3). To ensure an interpretable variance decomposition, we parameterized the total phenotypic variance using control parameters that partition it across shared and independent genetic, confounding, and noise components. The resulting allocation for the *A. thaliana* simulation is summarized in Supplementary Table S4. All simulated phenotypes were inverse-normal transformed prior to model fitting using the Van der Waerden procedure.

### Comparative study of multi- and univariate GWAS for metabolic phenotypes

To inspect the performance of TorchLIMIX on experimental data, we analyzed the dataset of Fusari et al. (7), which comprises metabolic traits (*i*.*e*., metabolite levels and enzyme activities) from primary metabolism measured across a diverse panel of 349 *A. thaliana* accessions from the HapMap panel (8; 7) in two experiments. The two experiments (Exp 1 and Exp 2) entail plants grown in a 10-h and 12-h photoperiod, with rosettes harvested at 37 and 28 days after sowing, respectively. Specifically, we selected the enzyme activities of the acid invertase (aINV) and the neutral invertase (nINV). Phenotypes were again inverse-normal transformed prior to model fitting using the Van der Waerden procedure. In the original study, Fusari et al. (7) performed univariate GWAS for each experiment separately. In contrast, our approach performs a joint multivariate LMM analysis, testing both experimental conditions simultaneously. Despite this methodological difference, the joint analysis is expected not only to recover the same significantly associated loci but also to reveal interaction effects that the univariate GWAS is unable to detect. Following Following Fusari et al. (7), we converted *p*-values to the LOD scale via LOD = −log_10_(*p*) and applied their threshold of LOD *≥* 3.0 as an initial filter to pinpoint loci identified by both uni- and multivariate GWAS. Fusari et al. applied a stricter threshold of LOD *≥* 4.8 for associations detected in only one experiment. We adopted the same threshold to identify additional SNPs not reported in the original study. SNPs exceeding the Bonferroni-corrected significance threshold are indicated separately. No minor allele frequency filtering was applied, consistent with the original study. SNPs were mapped to genes within a *±*10 kb window using the TAIR10 genome annotation, and genes were functionally categorized using the MapMan ontology (15) with mapping file (Ath_AGI_LOCUS_TAIR10_Aug2012) obtained from the PlaBiPD portal (https://www.plabipd.de).

## Results

To assess numerical equivalence, we compared TorchLIMIX and NumpyLIMIX across 10 simulations with four phenotypes and proportionality factors *η ∈* [*−*1, 1], as described in Supplementary Section 2. We evaluated differences in negative log marginal likelihoods (Δ LML), likelihood ratio test statistics (Δ LRT), and *p*-values under two initialization strategies for the covariance factor *L* of *C*_0_ = *LL*^*⊤*^: the default all-ones initialization (Supplementary Table S5) and the QR-based orthonormal initialization (Supplementary Table S6).

Under both strategies, mean absolute differences remained small across all metrics. For the default initialization, Δ LML values stayed below 6 *×* 10^*−*4^ and *p*-value discrepancies below 2 *×* 10^*−*4^ across all *η* values. For the QR-based initialization, typical Δ LML values were near 10^*−*4^–10^*−*3^, with occasional larger deviations at extreme *η* (up to 1.7 *×* 10^*−*2^ in Δ LML and 2.5 *×* 10^*−*3^ in Δ*p* at *η* = 1.0), reflecting differences in floating-point arithmetic across the underlying linear algebra backends (LAPACK/BLAS vs. cuBLAS/cuSOLVER) and operation ordering. Most importantly, the overlap in statistically significant loci (*p <* 5 *×* 10^*−*8^) remained 100% across all *η* values, all three likelihood ratio tests (LRT10, LRT20, LRT21), and both initialization strategies, confirming that TorchLIMIX reliably reproduces the results of the reference implementation.

### GPU-accelerated runtime performance across sample and phenotype dimensions

We compared NumpyLIMIX (CPU) and TorchLIMIX (GPU) for multivariate linear mixed model analyses. Both exploit the Kronecker structure of **C**_0_ ⊗ **K**_stable_ + **C**_1_ ⊗ **I** to avoid forming the full *np × np* covariance. Estimating **C**_0_ and **C**_1_ via L-BFGS-B costs *O*(*N*_iter_ *· k*^3^*r*^3^), where *k* = rank(**C**_0_) and *r* = rank(**G**). The null model is fitted once in *O*((*cp*)^3^) (*c*: covariates, *p*: phenotypes), and all null model dependent terms are precomputed and reused across all SNP tests. Each SNP test then costs *O*(*np* + (*cp* + *a*)^3^), where *a* is the number of columns in the alternative design matrix **A**_1_. Since SNP tests are independent, TorchLIMIX batches *b* SNPs along an extra dimension. The Cholesky pivot matrix depends only on the null model and is factorized once, so the *b* right-hand sides are concatenated into a single (*np × bk*) system and solved in one GPU kernel call. Chunks are processed sequentially, with intermediate results moved to CPU memory after each chunk to limit GPU memory usage. Hardware configurations are detailed in Supplementary Section 4. Benchmarks were performed using 18 CPU threads. Under these conditions, TorchLIMIX with GPU acceleration achieved substantial speedups over NumpyLIMIX. We benchmarked both implementations across sample sizes *n ∈ {*250, 300, 350, 400, 650, 1,300, 2,600, 5,200, 10,400, 20,800*}* and phenotype counts *p ∈ {*3, 4, 5, 6, 8, 10, 15*}* (Figure 1b, c;

Supplementary Table S7), reporting average runtimes over 10 independent runs. In the sample sweep (*p* = 4), speedups ranged from approximately 17*×* at *n* = 250 to 273*×* at *n* = 1,300, before decreasing at the largest sample sizes as GPU memory overhead grew (*≈* 25*×* at *n* = 20,800). In the phenotype sweep (*n* = 250), speedups increased from *≈* 17*×* (*p* = 4) to 85*×* (*p* = 8) as the computational cost of the CPU implementation scaled steeply with the number of phenotypes. Notably, TorchLIMIX scaled to configurations where NumpyLIMIX exceeded the 100-hour wall-clock budget (*p ≥* 10), completing in approximately 12 minutes for *p* = 10 and under 2.5 hours for *p* = 15.

### Improved p-value calibration through QR-based covariance initialization

The QR-based initialization of the Cholesky factor *L* substantially improved the statistical calibration of *p*-values. Under the default all-ones initialization, *p*-values exhibited notable inflation: the mean genomic inflation factor *λ* reached 1.222 *±* 0.208 and a Type I error rate of 0.0748 under the null hypothesis of the common effect test (*η* = *−*0.33, four phenotypes; Supplementary Figure 1a) and 1.583 *±* 0.295 and a Type I error rate of 0.0991 under the null hypothesis of the any vs. common effect test (*η* = 1.0; Supplementary Figure 1b). In contrast, the QR-based initialization yielded well-calibrated *p*-values across 1,000 simulated phenotypes, with genomic inflation factors of 1.004 *±* 0.011 and 1.011 *±* 0.018 for the common effect and any vs. common effect tests, respectively (Supplementary Figures 1c and 1d) and Type I error rates of 0.0499 and 0.0491. These results indicate that the QR-based initialization provides a more favorable optimization landscape for the covariance parameter estimation, effectively eliminating the systematic *p*-value inflation observed with the default strategy and ensuring reliable control of Type I error rates in multivariate association testing.

To investigate the source of this inflation, we examined the eigenvalue decomposition of the estimated covariance matrices across all 1,000 simulations (Supplementary Tables S8–S11). Under the default all-ones initialization, the genetic covariance matrix *C*_0_ consistently collapsed to an effective rank of exactly 1.000 *±* 0.000 in both the common effect (*η* = *−*0.33; Supplementary Table S8) and any vs. common effect (*η* = 1.0; Supplementary Table S9) null settings, despite being initialized at full rank. This rank collapse indicates convergence to a degenerate local optimum where *C*_0_ captures genetic covariance along only a single direction, inflating the likelihood ratio test statistics under the null. In contrast, the QR-based initialization preserved higher effective ranks of 2.088 *±* 0.333 and 1.674 *±* 0.268 under the common effect (Supplementary Table S10) and any vs. common effect (Supplementary Table S11) nulls, respectively. This higher-rank covariance structure enables a more accurate partitioning of variance between *C*_0_ and *C*_1_, yielding the well-calibrated *p*-values reported above.

### Comparison of univariate and multivariate GWAS

For *aINV*, our multivariate TorchLIMIX analysis confirmed 28 of 29 associated SNPs reported by Fusari et al. (7) (Figure 1d; Supplementary Figure 2a–e). The only SNP (*m87284*, chromosome 3) that was not recovered exhibited modest LOD scores in the original univariate analysis (LOD 3.4 and 3.1 in experiments 1 and 2, respectively) and did not reach significance in any test of our analysis. Four confirmed SNPs exceeded the Bonferroni threshold under LRT10 (LOD *>* 6.63). Beyond the confirmed loci, 14 additional significantly associated SNPs (LOD *>* 4.8) were identified, mapping to eight QTL regions, seven of which were not reported by Fusari et al., with three clustering within ~3 kb on chromosome 5 (Supplementary Figure 2a, e and Supplementary Data S3, Additional Files). The lead SNPs in two of the detected QTL regions approached the Bonferroni threshold (LOD 6.41 and 6.45). More details are provided in Supplementary Information, Section 7. Enrichment analysis at MapMan level 5 identified bin 2.2.1.3.3 (vacuolar invertases) as the only FDR-significant category (FDR = 1.6 × 10^−3^, fold enrichment = 82.65, Supplementary Table S12). At level 4, the parent bin 2.2.1.3 (invertases) was nominally significant alongside bins related to secondary metabolism and hormone signaling (Supplementary Table S12).

For *nINV*, all four SNPs found associated by Fusari et al. (7) were confirmed (Supplementary Figure 3a–e). One exceeded the Bonferroni threshold (*m196411* near *AT5G44560*, LOD = 7.11), while *m16610* near *AT1G27720* (LOD = 6.61) approached but did not reach Bonferroni significance (LOD = 6.63). Beyond the confirmed loci, 23 additional significantly associated SNPs (LOD *>* 4.8) were identified (Supplementary Data S3, Additional Files). One falls within an existing QTL region near *AT1G27720*, while the remaining 22 map to 18 additional QTL regions, most of which clustered near *AT1G27720* or *AT1G35530* and *AT1G35537* on chromosome 1 (Supplementary Figure 3a). As with *aINV*, the majority of additional signals were driven by the interaction and any effect tests rather than the common effect test. For *nINV*, no MapMan bins or GO terms reached FDR significance. Among nominally enriched MapMan categories, the vacuolar invertase bin (2.2.1.3.3, *p* = 0.016) was identified through the inclusion of *AT1G12240* (*VAC-INV*), the same locus underlying the strongest *aINV* associations. All significant loci (LOD *≥* 3.0) with gene annotations, cross-references to Fusari et al. SNPs, and functional classifications are listed in Supplementary Data S1 (*aINV*) and S2 (*nINV*) (Additional Files). No GO terms reached FDR significance for either trait. Complete MapMan and GO enrichment results are provided in Supplementary Data S4 and S6 for *aINV* and Supplementary Data S5 and S7 for *nINV* (Additional Files).

## Discussion

We demonstrated that the PyTorch implementation of LIMIX provides substantial computational acceleration of multivariate GWAS while maintaining numerical equivalence with the original framework. Notably, the scalability analysis was performed on the Horton dataset, which contains 173,220 SNPs after minor allele frequency filtering. Since TorchLIMIX’s batched GPU strategy parallelizes across SNPs, the speedup is expected to increase further for larger marker panels such as the *A. thaliana* 1001 Genomes dataset (1). Moreover, a revised initialization of the genetic covariance matrix prevents rank-1 collapse during optimization, yielding better-calibrated *p*-values and Type-I error control. The novel G*×*E-driven loci identified here for both aINV and nINV, which were not detected in the original univariate GWAS, underscore the practical value of scalable and numerically robust multivariate implementations.

## Author contribution

Bibiana M. Horn (Conceptualization [lead], Methodology [lead], Software [lead], Formal Analysis [lead], Visualization [lead], Investigation [lead], Writing—original draft [lead], Writing—review & editing [lead]), Zoran Nikoloski (Supervision [supporting], Writing—review & editing [supporting]), and Christoph Lippert (Conceptualization [supporting], Methodology [supporting], Supervision [lead], Writing—review & editing [supporting])

## Supplementary Materials

Supplementary figures and tables are provided in a separate Supplementary PDF. Additional data files (Supplementary Data S1– S7) are provided as Excel files.

## Data availability

This study uses previously published experimental data (7). Code for the *in silico* phenotype simulations is available at https://github.com/bi-horn/torchLIMIX.

## Acknowledgments

Use of AI tools

Claude (Anthropic) was used as an aid to polish written text, formatting tables, refactor computational scripts, and draft documentation for code functions. All outputs were critically reviewed, verified, and edited by the authors, who take full responsibility for the accuracy of the final text and code.

