## Supplementary PDF for "TorchLIMIX: GPU-accelerated multivariate genome-wide association studies"

### Supplementary Material for: "TorchLIMIX: GPU-accelerated multivariate genome-wide association studies"

#### Abstract

This supplementary material provides additional detail on the implementation, validation, and application of TorchLIMIX. We describe the covariance parameterization and structured hypothesis testing framework, followed by the phenotype simulation design adapted from Casale et al. (2017). Validation results include numerical equivalence of likelihood values between the NumPy and PyTorch implementations, scalability benchmarks, $p$-value calibration under default and refactored genetic covariance initialization, and effective rank comparison for both initialization strategies. Furthermore, per-chromosome Manhattan plots and MapMan bin enrichment analyses are presented for acid and neutral invertase activity, based on data from *A. thaliana* experiments, obtained from Fusari et al. (2017).

#### Covariance implementation and structured hypothesis testing

To compute the variance-normalized genetic relatedness matrix $K_{\text{stable}}$ referenced in the main text, we first calculate the kinship matrix $K = GG^\top/d$, where $G \in \mathbb{R}^{n\times d}$ is the standardized genotype matrix. An economic eigendecomposition yields $K_{\text{stable}} = QSQ^\top$ and the low-rank factorization $G_{\text{stable}} = Q\sqrt{S}$. TorchLIMIX applies min-max scaling to the genetic relatedness matrix, consistent with the original LIMIX framework (3). When scaling is used, the covariance parameter $C_0$ is estimated in the normalized space and rescaled by $G_{\text{norm}}^{-2}$ to recover the original parameterization.

The genetic covariance is factorized as $C_0 = L_0 L_0^\top$ with $L_0 \in \mathbb{R}^{p\times r}$, where $L_0$ is a dense, unconstrained factor. For $L_0$, TorchLIMIX supports two initialization strategies:

1. **QR-based (default):** $L_0$ is set to the orthonormal factor $Q$ obtained from the QR decomposition of a random matrix. Since $Q^\top Q = I_r$, the initial covariance $C_0 = QQ^\top$ has all nonzero eigenvalues equal to unity, yielding a well-conditioned starting point regardless of rank.
2. **Original LIMIX:** all entries of $L_0$ are set to one, matching the initialization used in the original LIMIX framework (3).

The error covariance is parameterized as $C_1 = L_1 L_1^\top + \varepsilon I_p$ with $\varepsilon > 0$ for numerical stability, where $L_1$ is lower-triangular with logarithmically parameterized diagonal elements to ensure positivity. The covariance matrices $C_0$ and $C_1$ are optimized using analytical gradients as in `NumPyLIMIX` (3).

After convergence, fixed effects ($B_{\text{cov}}$, $B_{\text{snp}}$) are obtained in closed form via generalized least squares. The trait-design matrices $A_{\text{cov}} \in \mathbb{R}^{p\times \cdot}$ and $A_{\text{snp}} \in \mathbb{R}^{p\times \cdot}$ specify which phenotypes are influenced by covariates and the genetic variant, respectively. Their construction and the degrees of freedom for each hypothesis test are detailed in Table S1. Statistical significance is determined by comparing the resulting log-likelihood ratio test statistics against a $\chi^2$ distribution with degrees of freedom equal to the difference in free parameters between the alternative and null models.

**Table S1** Extended null and alternative hypotheses, design matrices, and degrees of freedom (DoF) for multivariate hypothesis tests with $p$ phenotypes.

| Test Type | Null Hypothesis ($H_0$) | Alternative ($H_A$) | Design Matrices | DoF |
|---|---|---|---|---|
| Common Effect | $\beta_1 = \cdots = \beta_p = 0$ | $\beta_1 = \cdots = \beta_p \neq 0$ | $A_0 = 0_p$<br>$A_1 = \mathbf{1}_p$ | 1 |
| Any Effect | $\beta_1 = \cdots = \beta_p = 0$ | $\exists\, p : \beta_p \neq 0$ | $A_0 = 0_p$<br>$A_1 = I_p$ | $p$ |
| Specific Effect | $\beta_p = 0$ | $\beta_p \neq 0$ | $A_0 = 0_p$<br>$A_1 = \mathbf{e}_p$ | 1 |
| Specific vs. Common | $\beta_p = \bar{\beta}$ | $\beta_p \neq \bar{\beta}$ | $A_0 = \mathbf{1}_p$<br>$A_1 = [\mathbf{1}_p\ \mathbf{e}_p]$ | 1 |
| Any Specific vs. Common | $\beta_1 = \cdots = \beta_p$ | $\exists\, p : \beta_p \neq \bar{\beta}$ | $A_0 = \mathbf{1}_p$<br>$A_1 = I_p$ | $p - 1$ |

*Note:* $I_p$ denotes the $p \times p$ identity matrix; $\mathbf{1}_p$ and $0_p$ are vectors of ones and zeros; $\mathbf{e}_p$ is the unit vector with a 1 at position $p$; $\bar{\beta}$ denotes the common effect across traits.

#### In silico phenotype simulation model

For a randomly sampled set of $n$ individuals with genotype matrix $\mathbf{G}$ of dimensions $n \times s$ ($s$ is the number of SNPs), we generate $p$ phenotypes $\mathbf{Y}$ of dimensions $n \times p$ as the sum of multiple components, following the structure proposed by Casale et al. (1):

$$Y = S + G + H + \Psi \ , \tag{1}$$

where $S$ represents the causal SNP effect, $G = G^{(s)} + G^{(i)}$ denotes the polygenic background effects decomposed into shared and independent components, $H = H^{(s)} + H^{(i)}$ captures hidden confounding effects similarly decomposed, and $\Psi$ represents independent and identically distributed (i.i.d.) noise.

The causal SNP effect $S$ is controlled by a rescaling parameter $\eta$ that governs the proportional change in effect sizes across phenotypes and is computed as:

$$\mathbf{S} = \mathbf{G}_{\text{causal}}\mathbf{b}\,\mathbf{v}^{\top}, \qquad \mathbf{v} = (1, \eta, \ldots, \eta) \in \mathbb{R}^{p}, \tag{2}$$

where $\mathbf{G}_{\text{causal}}$ is the genotype vector for the causal variant, the effect size $b$ is drawn from $\{-1, +1\}$, and $\eta$ is the rescaling factor controlling the proportional change in effect sizes across phenotypes (1). The total effect variance splits into persistent and rescaling fractions defined as $f_{\text{persistent}} = (1 + (P-1)\eta)^2 / \big[P(1 + (P-1)\eta^2)\big]$ and $f_{\text{rescaling}} = (1-\eta)^2(P-1) / \big[P(1 + (P-1)\eta^2)\big]$, which sum to one by construction. Supplementary Table S2 reports both fractions for four phenotypes across a range of $\eta$ values. For $p = 4$ phenotypes, $\eta = -0.33$ maximizes the phenotype-specific variance, whereas $\eta = 1.0$ produces fully persistent effects shared across all phenotypes. The polygenic background effects, hidden confounding effects, and noise were sampled from matrix normal distributions as described in Supplementary Table S3 following the description by Casale et al. (1). To simulate interpretable variance components, we decompose the total phenotypic variance using control parameters as summarized in Supplementary Table S4. Each raw component is then rescaled to match the target variance fractions:

$$X_{\text{final}} = X_{\text{raw}} \cdot \sqrt{v_{\text{target}} \,/\, \mathrm{Var}[\mathrm{vec}(X_{\text{raw}})]}\,, \tag{3}$$

where $X$ stands for any of $S$, $G^{(s)}$, $G^{(i)}$, $H^{(s)}$, $H^{(i)}$, or $\Psi$.

**Table S2** Variance fractions of persistent and rescaling effects across different $\eta$ values for $p = 4$ phenotypes

| $\eta$ | $f_{\text{persistent}}$ | $f_{\text{rescaling}}$ | Sum |
|---|---|---|---|
| $-1.0$ | 0.2500 | 0.7500 | 1.0000 |
| $-0.8$ | 0.1678 | 0.8322 | 1.0000 |
| $-0.6$ | 0.0769 | 0.9231 | 1.0000 |
| $-0.4$ | 0.0068 | 0.9932 | 1.0000 |
| $-0.33$ | 0.0000 | 1.0000 | 1.0000 |
| $-0.2$ | 0.0357 | 0.9643 | 1.0000 |
| $0.0$ | 0.2500 | 0.7500 | 1.0000 |
| $0.2$ | 0.5714 | 0.4286 | 1.0000 |
| $0.4$ | 0.8176 | 0.1824 | 1.0000 |
| $0.6$ | 0.9423 | 0.0577 | 1.0000 |
| $0.8$ | 0.9897 | 0.0103 | 1.0000 |
| $1.0$ | 1.0000 | 0.0000 | 1.0000 |

**Table S3** Matrix normal distributions for simulating polygenic background, hidden confounding, and noise effects.

| Component | Distribution | Description |
|---|---|---|
| **Polygenic Background Effects ($G$)** | | |
| $G^{(s)}$ | $\mathcal{MN}(\mathbf{0},\, \mathbf{K},\, a_G^2\, \mathbf{1}_P \mathbf{1}_P^\top)$ | Shared polygenic background |
| $G^{(i)}$ | $\mathcal{MN}(\mathbf{0},\, \mathbf{K},\, c_G^2\, \mathbf{I}_P)$ | Independent polygenic background |
| **Hidden Confounding Effects ($H$)** | | |
| $H^{(s)}$ | $\mathcal{MN}(\mathbf{0},\, \mathbf{M}\mathbf{M}^\top,\, a_H^2\, \mathbf{1}_P \mathbf{1}_P^\top)$ | Shared hidden confounding |
| $H^{(i)}$ | $\mathcal{MN}(\mathbf{0},\, \mathbf{M}\mathbf{M}^\top,\, c_H^2\, \mathbf{I}_P)$ | Independent hidden confounding |
| **Noise Component ($\Psi$)** | | |
| $\Psi_{ij}$ | $\mathcal{N}(0,\, \sigma^2)$ | i.i.d. noise per entry |

*Notation.* $\mathcal{MN}$ denotes the matrix normal distribution; $\mathbf{K}$ is the kinship matrix; $\mathbf{M} \in \mathbb{R}^{N \times L}$ has entries $M_{ij} \sim \mathcal{N}(0, 1)$, where $L$ is the number of hidden confounders. *Parameter sampling.* For each replicate, $a_G = \sqrt{\alpha_G}$ and $c_G = \sqrt{\gamma_G}$ with $\alpha_G,\ \gamma_G \sim \mathrm{U}(0, 1)$; $a_H$ and $c_H$ are obtained analogously.

**Table S4** Variance allocation across different components for phenotype simulation. Selected control parameters for the simulation: $v_s = 0.15$, $v_{\text{bg}} = 0.50$, $\phi = 0.6$, $\gamma = 0.4$.

| Component | Symbol | Description | Variance formula | Resulting variances |
|---|---|---|---|---|
| Region | S | Causal variant | $v_s$ | 0.150 |
| Shared Background | $G^{(s)}$ | Shared polygenic background | $\phi \cdot v_{\text{bg}}$ | 0.300 |
| Independent Background | $G^{(i)}$ | Independent polygenic background | $(1 - \phi) \cdot v_{\text{bg}}$ | 0.200 |
| Shared Hidden | $H^{(s)}$ | Shared hidden confounding | $\phi \cdot \gamma \cdot (1 - v_{\text{bg}} - v_s)$ | 0.084 |
| Independent Hidden | $H^{(i)}$ | Independent hidden confounding | $(1 - \phi) \cdot \gamma \cdot (1 - v_{\text{bg}} - v_s)$ | 0.056 |
| Independent Noise | $\Psi$ | i.i.d. noise | $(1 - \gamma) \cdot (1 - v_{\text{bg}} - v_s)$ | 0.210 |
| **Total** | | | | **1.000** |

*Note:* $v_s$ = causal variant variance; $\phi$ = proportion shared vs. independent; $\gamma$ = proportion hidden confounding vs. noise; $v_{\text{bg}}$ = polygenic background variance.

#### Numerical equivalence of likelihood values between NumPy and Torch pipeline implementation

Supplementary Tables S5 and S6 report the likelihood agreement between NumPyLIMIX and TorchLIMIX across proportionality factor $\eta \in [-1, 1]$ for the default and orthogonal initialization of $\mathbf{C}_0$, respectively.

**Table S5** Likelihood agreement of NumPyLIMIX and TorchLIMIX across four simulated phenotypes scaled by proportionality factors $\eta$ for the default initialization of the genetic covariance matrix $\mathbf{C}_0$

| $\eta$ | Test | $\overline{\lvert\Delta \mathrm{LML}\rvert}$ | $\overline{\lvert\Delta \mathrm{LRT}\rvert}$ | $\overline{\lvert\Delta p\rvert}$ | Significant NumPy | Significant Torch | Overlap |
|---|---|---|---|---|---|---|---|
| $-1.0$ | LRT10 | 1.3e-4 ± 1.5e-4 | 2.6e-4 ± 3.0e-4 | 7.7e-5 ± 8.8e-5 | 3 | 3 | 1.00 |
| | LRT20 | 2.6e-4 ± 3.0e-4 | 5.3e-4 ± 6.0e-4 | 4.5e-5 ± 5.2e-5 | 142 | 142 | 1.00 |
| | LRT21 | 2.6e-4 ± 3.0e-4 | 4.5e-4 ± 5.0e-4 | 4.5e-5 ± 5.1e-5 | 111 | 111 | 1.00 |
| $-0.8$ | LRT10 | 1.6e-4 ± 1.8e-4 | 3.0e-4 ± 3.6e-4 | 9.6e-5 ± 1.2e-4 | 1 | 1 | 1.00 |
| | LRT20 | 3.2e-4 ± 3.6e-4 | 6.3e-4 ± 7.2e-4 | 5.8e-5 ± 6.9e-5 | 137 | 137 | 1.00 |
| | LRT21 | 3.2e-4 ± 3.6e-4 | 5.5e-4 ± 6.1e-4 | 5.8e-5 ± 6.8e-5 | 120 | 120 | 1.00 |
| $-0.6$ | LRT10 | 8.3e-5 ± 8.2e-5 | 1.4e-4 ± 1.4e-4 | 4.2e-5 ± 3.8e-5 | 4 | 4 | 1.00 |
| | LRT20 | 1.4e-4 ± 1.3e-4 | 2.7e-4 ± 2.6e-4 | 2.3e-5 ± 2.3e-5 | 108 | 108 | 1.00 |
| | LRT21 | 1.4e-4 ± 1.3e-4 | 2.1e-4 ± 2.1e-4 | 2.2e-5 ± 2.2e-5 | 126 | 126 | 1.00 |
| $-0.4$ | LRT10 | 1.4e-4 ± 1.6e-4 | 2.7e-4 ± 3.0e-4 | 8.7e-5 ± 9.7e-5 | 2 | 2 | 1.00 |
| | LRT20 | 3.2e-4 ± 4.0e-4 | 6.4e-4 ± 8.0e-4 | 5.7e-5 ± 7.3e-5 | 105 | 105 | 1.00 |
| | LRT21 | 3.2e-4 ± 4.0e-4 | 5.6e-4 ± 7.3e-4 | 5.9e-5 ± 7.9e-5 | 130 | 130 | 1.00 |
| $-0.2$ | LRT10 | 1.5e-4 ± 2.5e-4 | 3.0e-4 ± 4.9e-4 | 9.9e-5 ± 1.7e-4 | 2 | 2 | 1.00 |
| | LRT20 | 3.2e-4 ± 5.3e-4 | 6.4e-4 ± 1.1e-3 | 5.7e-5 ± 9.6e-5 | 99 | 99 | 1.00 |
| | LRT21 | 3.2e-4 ± 5.3e-4 | 5.6e-4 ± 9.4e-4 | 5.7e-5 ± 9.7e-5 | 110 | 110 | 1.00 |
| 0.0 | LRT10 | 1.3e-4 ± 1.8e-4 | 2.5e-4 ± 3.6e-4 | 8.1e-5 ± 1.2e-4 | 4 | 4 | 1.00 |
| | LRT20 | 2.9e-4 ± 4.7e-4 | 5.7e-4 ± 9.3e-4 | 4.9e-5 ± 8.2e-5 | 70 | 70 | 1.00 |
| | LRT21 | 2.9e-4 ± 4.7e-4 | 4.9e-4 ± 8.3e-4 | 4.9e-5 ± 8.6e-5 | 68 | 68 | 1.00 |
| 0.2 | LRT10 | 1.3e-4 ± 1.7e-4 | 2.0e-4 ± 2.2e-4 | 6.5e-5 ± 7.3e-5 | 10 | 10 | 1.00 |
| | LRT20 | 2.4e-4 ± 3.0e-4 | 4.6e-4 ± 5.5e-4 | 3.9e-5 ± 4.7e-5 | 49 | 49 | 1.00 |
| | LRT21 | 2.4e-4 ± 3.0e-4 | 4.0e-4 ± 4.9e-4 | 4.0e-5 ± 4.9e-5 | 29 | 29 | 1.00 |
| 0.4 | LRT10 | 1.2e-4 ± 1.4e-4 | 2.4e-4 ± 2.8e-4 | 7.9e-5 ± 9.6e-5 | 16 | 16 | 1.00 |
| | LRT20 | 2.3e-4 ± 2.6e-4 | 4.6e-4 ± 5.2e-4 | 4.1e-5 ± 4.7e-5 | 37 | 37 | 1.00 |
| | LRT21 | 2.3e-4 ± 2.6e-4 | 3.9e-4 ± 4.4e-4 | 4.0e-5 ± 4.6e-5 | 14 | 14 | 1.00 |
| 0.6 | LRT10 | 9.8e-5 ± 1.1e-4 | 1.9e-4 ± 2.3e-4 | 6.3e-5 ± 7.3e-5 | 16 | 16 | 1.00 |
| | LRT20 | 2.0e-4 ± 2.5e-4 | 4.1e-4 ± 5.0e-4 | 3.4e-5 ± 4.0e-5 | 69 | 69 | 1.00 |
| | LRT21 | 2.0e-4 ± 2.5e-4 | 3.6e-4 ± 4.6e-4 | 3.3e-5 ± 4.0e-5 | 62 | 62 | 1.00 |
| 0.8 | LRT10 | 2.4e-4 ± 4.2e-4 | 4.9e-4 ± 8.4e-4 | 1.5e-4 ± 2.6e-4 | 25 | 25 | 1.00 |
| | LRT20 | 4.9e-4 ± 8.3e-4 | 9.8e-4 ± 1.6e-3 | 8.4e-5 ± 1.4e-4 | 43 | 43 | 1.00 |
| | LRT21 | 4.9e-4 ± 8.3e-4 | 8.0e-4 ± 1.4e-3 | 7.9e-5 ± 1.4e-4 | 23 | 23 | 1.00 |
| 1.0 | LRT10 | 4.0e-4 ± 1.1e-3 | 6.3e-4 ± 1.7e-3 | 1.9e-4 ± 5.0e-4 | 24 | 24 | 1.00 |
| | LRT20 | 5.5e-4 ± 1.4e-3 | 1.0e-3 ± 2.6e-3 | 9.3e-5 ± 2.4e-4 | 29 | 29 | 1.00 |
| | LRT21 | 5.5e-4 ± 1.4e-3 | 7.7e-4 ± 2.0e-3 | 8.4e-5 ± 2.2e-4 | 24 | 24 | 1.00 |

*Note:* Likelihood ratio tests (LRTs) — LRT10: common effect vs. null; LRT20: any effect vs. null; LRT21: any vs. common effect (interaction). $\overline{\lvert\Delta\cdot\rvert}$: for each replicate, the absolute difference between implementations is computed per SNP and averaged; the table reports mean ± std of these averages across $n{=}10$ replicates. Significant NumPy / Significant Torch: cumulative significant loci ($p < 2.89 \times 10^{-7}$, Bonferroni correction for 173,220 tests) for NumPyLIMIX / TorchLIMIX. Overlap: fraction of SNPs with identical significance call.

**Table S6** Likelihood agreement of NumPyLIMIX and TorchLIMIX across four simulated phenotypes scaled by proportionality factor $\eta$ for the refactored initialization of the genetic covariance matrix $\mathbf{C}_0$

| $\eta$ | Test | $\overline{\lvert\Delta \mathrm{LML}\rvert}$ | $\overline{\lvert\Delta \mathrm{LRT}\rvert}$ | $\overline{\lvert\Delta p\rvert}$ | Significant NumPy | Significant Torch | Overlap |
|---|---|---|---|---|---|---|---|
| $-1.0$ | LRT10 | 3.4e-4 ± 4.4e-4 | 4.3e-4 ± 3.2e-4 | 1.5e-4 ± 1.1e-4 | 0 | 0 | 1.00 |
| | LRT20 | 5.2e-4 ± 4.7e-4 | 8.7e-4 ± 5.9e-4 | 9.6e-5 ± 6.5e-5 | 70 | 70 | 1.00 |
| | LRT21 | 5.2e-4 ± 4.7e-4 | 7.1e-4 ± 4.7e-4 | 9.3e-5 ± 6.2e-5 | 70 | 70 | 1.00 |
| $-0.8$ | LRT10 | 4.3e-4 ± 5.5e-4 | 5.0e-4 ± 5.6e-4 | 1.8e-4 ± 2.0e-4 | 0 | 0 | 1.00 |
| | LRT20 | 6.6e-4 ± 6.9e-4 | 1.1e-3 ± 1.1e-3 | 1.2e-4 ± 1.2e-4 | 76 | 76 | 1.00 |
| | LRT21 | 6.6e-4 ± 6.9e-4 | 9.1e-4 ± 9.3e-4 | 1.2e-4 ± 1.2e-4 | 84 | 84 | 1.00 |
| $-0.6$ | LRT10 | 3.3e-4 ± 3.5e-4 | 5.0e-4 ± 4.4e-4 | 1.7e-4 ± 1.6e-4 | 0 | 0 | 1.00 |
| | LRT20 | 5.1e-4 ± 4.2e-4 | 9.2e-4 ± 7.1e-4 | 1.0e-4 ± 7.9e-5 | 79 | 79 | 1.00 |
| | LRT21 | 5.1e-4 ± 4.2e-4 | 7.1e-4 ± 5.3e-4 | 9.3e-5 ± 7.1e-5 | 98 | 98 | 1.00 |
| $-0.4$ | LRT10 | 3.9e-4 ± 4.3e-4 | 6.3e-4 ± 6.1e-4 | 2.2e-4 ± 2.2e-4 | 0 | 0 | 1.00 |
| | LRT20 | 6.5e-4 ± 6.2e-4 | 1.2e-3 ± 1.1e-3 | 1.4e-4 ± 1.2e-4 | 75 | 75 | 1.00 |
| | LRT21 | 6.5e-4 ± 6.2e-4 | 1.0e-3 ± 8.9e-4 | 1.3e-4 ± 1.2e-4 | 96 | 96 | 1.00 |
| $-0.2$ | LRT10 | 2.0e-3 ± 4.4e-3 | 5.7e-4 ± 4.6e-4 | 2.0e-4 ± 1.6e-4 | 0 | 0 | 1.00 |
| | LRT20 | 2.2e-3 ± 4.3e-3 | 9.8e-4 ± 6.9e-4 | 1.1e-4 ± 7.6e-5 | 69 | 69 | 1.00 |
| | LRT21 | 2.2e-3 ± 4.3e-3 | 7.6e-4 ± 5.0e-4 | 1.0e-4 ± 6.7e-5 | 82 | 82 | 1.00 |
| $0.0$ | LRT10 | 5.9e-4 ± 7.0e-4 | 4.9e-4 ± 3.9e-4 | 1.7e-4 ± 1.4e-4 | 1 | 1 | 1.00 |
| | LRT20 | 7.7e-4 ± 7.1e-4 | 1.1e-3 ± 7.9e-4 | 1.2e-4 ± 8.7e-5 | 43 | 43 | 1.00 |
| | LRT21 | 7.7e-4 ± 7.1e-4 | 9.2e-4 ± 6.6e-4 | 1.2e-4 ± 8.7e-5 | 41 | 41 | 1.00 |
| $0.2$ | LRT10 | 2.4e-4 ± 2.3e-4 | 4.3e-4 ± 4.0e-4 | 1.5e-4 ± 1.4e-4 | 8 | 8 | 1.00 |
| | LRT20 | 4.2e-4 ± 3.7e-4 | 8.1e-4 ± 7.3e-4 | 8.9e-5 ± 8.2e-5 | 33 | 33 | 1.00 |
| | LRT21 | 4.2e-4 ± 3.7e-4 | 6.3e-4 ± 6.0e-4 | 8.4e-5 ± 7.9e-5 | 18 | 18 | 1.00 |
| $0.4$ | LRT10 | 2.8e-4 ± 3.4e-4 | 5.3e-4 ± 6.5e-4 | 1.9e-4 ± 2.3e-4 | 13 | 13 | 1.00 |
| | LRT20 | 4.7e-4 ± 4.7e-4 | 9.3e-4 ± 9.2e-4 | 1.0e-4 ± 1.0e-4 | 22 | 22 | 1.00 |
| | LRT21 | 4.7e-4 ± 4.7e-4 | 6.8e-4 ± 6.4e-4 | 8.9e-5 ± 8.4e-5 | 3 | 3 | 1.00 |
| $0.6$ | LRT10 | 2.4e-4 ± 2.7e-4 | 4.6e-4 ± 5.2e-4 | 1.6e-4 ± 1.8e-4 | 15 | 15 | 1.00 |
| | LRT20 | 4.1e-4 ± 4.1e-4 | 8.1e-4 ± 8.1e-4 | 8.9e-5 ± 8.9e-5 | 14 | 14 | 1.00 |
| | LRT21 | 4.1e-4 ± 4.1e-4 | 6.1e-4 ± 5.6e-4 | 8.0e-5 ± 7.4e-5 | 1 | 1 | 1.00 |
| $0.8$ | LRT10 | 8.6e-3 ± 2.6e-2 | 3.7e-3 ± 1.0e-2 | 1.3e-3 ± 3.6e-3 | 17 | 17 | 1.00 |
| | LRT20 | 8.8e-3 ± 2.6e-2 | 5.9e-3 ± 1.6e-2 | 6.5e-4 ± 1.8e-3 | 12 | 12 | 1.00 |
| | LRT21 | 8.8e-3 ± 2.6e-2 | 4.2e-3 ± 1.1e-2 | 5.6e-4 ± 1.5e-3 | 0 | 0 | 1.00 |
| $1.0$ | LRT10 | 1.7e-2 ± 3.8e-2 | 7.1e-3 ± 1.5e-2 | 2.5e-3 ± 5.3e-3 | 10 | 10 | 1.00 |
| | LRT20 | 1.7e-2 ± 3.8e-2 | 1.1e-2 ± 2.3e-2 | 1.2e-3 ± 2.6e-3 | 6 | 6 | 1.00 |
| | LRT21 | 1.7e-2 ± 3.8e-2 | 7.8e-3 ± 1.6e-2 | 1.0e-3 ± 2.2e-3 | 0 | 0 | 1.00 |

*Note:* Likelihood ratio tests (LRTs) — LRT10: common effect vs. null; LRT20: any effect vs. null; LRT21: any vs. common effect (interaction). $\overline{\lvert\Delta \cdot\rvert}$: for each replicate, the absolute difference between implementations is computed per SNP and averaged; the table reports mean ± std of these averages across $n{=}10$ replicates (except $\eta{=}1.0$ where $n{=}5$). Significant NumPy / Significant Torch: cumulative significant loci ($p < 2.89 \times 10^{-7}$, Bonferroni correction for 173,220 tests) for NumPyLIMIX / TorchLIMIX. Overlap: fraction of SNPs with identical significance call.

#### Scalability analysis

Table S7 reports the runtime comparison between NumpyLIMIX and TorchLIMIX across varying numbers of samples and phenotypes. Runtimes are reported as mean ± SD over 10 independent runs with $\eta = 0.5$ and $v_s = 0.05$, chosen to reflect a low GxE phenotype setting. Each run uses a different simulated phenotype, with both implementations evaluated on the same phenotype within each run to ensure a fair comparison. The HapMap panel dataset contains $n = 1{,}307$ samples (2). For configurations exceeding this size ($n \in \{2{,}600,\ 5{,}200,\ 10{,}400,\ 20{,}800\}$), the genome dataset was stacked to generate the required number of observations.

Both implementations use 18 CPU threads and 100 GB of system memory. The CPU-only configuration used an AMD EPYC 7742 64-Core Processor (2.25 GHz base), while the GPU configuration used a single NVIDIA A100-SXM4-40GB on a node equipped with an AMD EPYC 7662 64-Core Processor (2.0 GHz base). This CPU difference is expected to have minimal impact, as TorchLIMIX offloads the computationally intensive linear algebra to the GPU, making its runtime largely dependent on GPU rather than host CPU performance.

We imposed a total wall-clock budget of 100 hours per configuration. The $p = 10$ configuration for NumpyLIMIX completed only 4 of 10 runs within this budget; the $p = 15$ configuration did not complete any run. Therefore, missing entries for NumpyLIMIX in the phenotype sweep indicate configurations that exceeded this budget.

The number of L-BFGS-B iterations required for null model convergence is reported alongside each runtime, as this varies with the likelihood surface of each configuration and explains why runtimes do not always increase monotonically with problem size. In the sample sweep, both implementations converge in the same number of iterations, since only the data size changes while the parameter space remains fixed. In the phenotype sweep, however, iteration counts increasingly diverge for $p \geq 5$. This is expected because the number of covariance parameters grows with $p$, producing a higher dimensional optimization landscape where small numerical differences between backends, such as different BLAS and LAPACK implementations, rounding behavior in matrix operations, and nondeterministic summation order under multithreading, accumulate across iterations and lead to different optimizer trajectories. While the likelihood validation (Section 3) used a single thread with deterministic arithmetic, that setting is considerably slower and was therefore not used for benchmarking.

**Table S7** Runtime comparison between NumPyLIMIX and TorchLIMIX on a single NVIDIA A100 GPU and an AMD EPYC 64-Core CPU (18 threads, 100 GB RAM). **(A)** Sample sweep with number of phenotypes fixed at 4. **(B)** Phenotype sweep with number of samples fixed at 250. Runtimes are reported as mean ± standard deviation over 10 independent runs.

| | NumPyLIMIX | | TorchLIMIX | | |
|---|---|---|---|---|---|
| $n/p$ | Runtime [min] | Iter. | Runtime [min] | Iter. | Speedup |
| *(A) Sample sweep (number of phenotypes = 4)* | | | | | |
| 250 | 7.03 ± 1.23 | 71 | 0.42 ± 0.21 | 71 | 16.7 |
| 300 | 11.25 ± 2.94 | 50 | 0.40 ± 0.15 | 50 | 28.4 |
| 350 | 14.11 ± 3.24 | 39 | 0.31 ± 0.12 | 39 | 44.8 |
| 400 | 20.45 ± 0.99 | 43 | 0.29 ± 0.03 | 43 | 71.6 |
| 650 | 55.40 ± 3.15 | 33 | 0.39 ± 0.02 | 33 | 143.7 |
| 1300 | 240.70 ± 10.61 | 25 | 0.88 ± 0.04 | 25 | 272.8 |
| 2600 | 246.33 ± 51.02 | 19 | 1.06 ± 0.03 | 19 | 233.4 |
| 5200 | 226.71 ± 8.03 | 17 | 3.51 ± 0.44 | 17 | 64.6 |
| 10400 | 229.33 ± 9.67 | 18 | 10.46 ± 0.12 | 18 | 21.9 |
| 20800 | 511.12 ± 181.62 | 21 | 20.55 ± 0.81 | 21 | 24.9 |
| *(B) Phenotype sweep (number of samples = 250)* | | | | | |
| 3 | 4.96 ± 0.34 | 37 | 0.22 ± 0.05 | 37 | 22.6 |
| 4 | 7.03 ± 1.23 | 71 | 0.42 ± 0.21 | 71 | 16.7 |
| 5 | 21.64 ± 9.28 | 132 | 0.80 ± 0.36 | 147 | 26.9 |
| 6 | 68.62 ± 31.30 | 229 | 1.66 ± 0.92 | 236 | 41.2 |
| 8 | 411.05 ± 269.37 | 398 | 4.82 ± 2.04 | 349 | 85.3 |
| 10 | – | – | 12.22 ± 6.14 | 381 | – |
| 15 | – | – | 136.57 ± 37.85 | 1282 | – |

*Note:* All runtimes are averaged over 10 runs. A total wall-clock budget of 100 hours was imposed per configuration; missing entries indicate that the budget was exceeded before all runs completed. Speedup = Runtime NumPyLIMIX / Runtime TorchLIMIX. Iter. = median number of L-BFGS-B iterations (null-model fitting) across runs.

#### Calibration of hypothesis test p-values

Supplementary Figure 1 shows quantile–quantile (QQ) plots of $p$-values under the null hypothesis (1,000 repetitions) for the common effect test (LRT10; panels a, c), and the any vs. common effect test (LRT21; panels b, d).

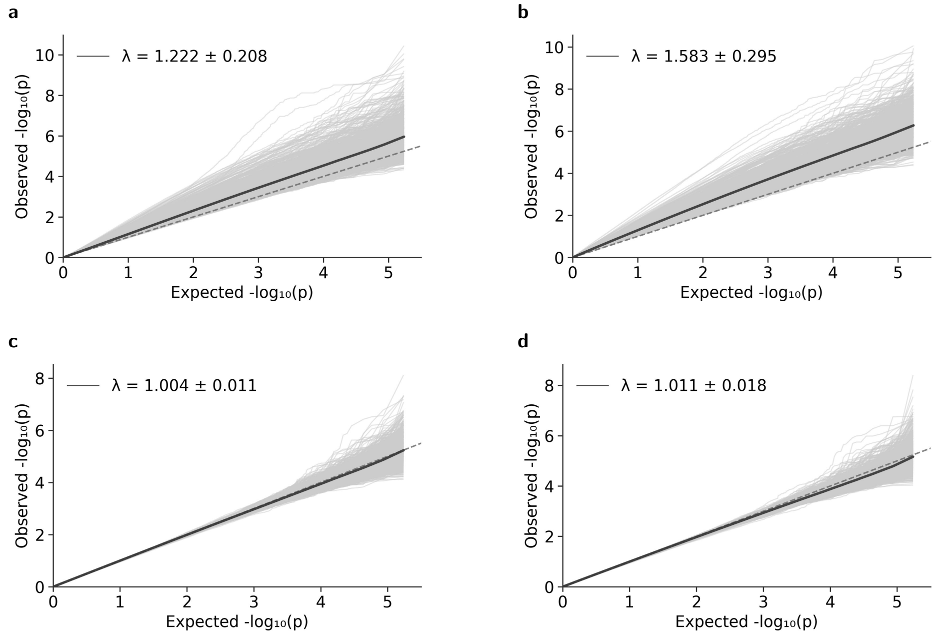


**Figure 1** Comparison of p-value calibration for TorchLIMIX with default and refactored initialization of the genetic covariance matrix $C_0$ (1000 repetitions under the null hypothesis of the respective hypothesis test). **(a)** QQ plot of $p$-values from the common effect test (LRT10) with default initialization of $C_0$. **(b)** QQ plot of $p$-values from the any vs. common effect test (LRT21) with default initialization of $C_0$. **(c)** QQ plot of $p$-values from the common effect test (LRT10) with refactored, QR-based initialization of $C_0$. **(d)** QQ plot of $p$-values from the any vs. common effect test (LRT21) with refactored, QR-based initialization of $C_0$. Mean genomic inflation factors $\lambda$ are shown in each panel.

#### Effective rank comparison for different genetic covariance initializations

Tables S8–S9 summarize the eigenvalue decomposition of the estimated genetic ($C_0$) and error ($C_1$) covariance matrices under different null hypotheses for the default initialization, while Tables S10–S11 present the corresponding results for the QR-based initialization. Each table reports the mean and standard deviation of the eigenvalues $\lambda_1$ to $\lambda_4$ and the effective rank across n = 1000 simulation repetitions. The effective rank is computed as the participation ratio $e_{\text{rank}} = \left(\sum_i \lambda_i\right)^2 / \sum_i \lambda_i^2$, which equals the matrix dimension when all eigenvalues are equal and reduces to one when a single eigenvalue dominates. These diagnostics serve to characterize the rank structure recovered by the optimizer and to verify that the covariance estimates are consistent with the constraints imposed by each null hypothesis.

**Table S8** Eigenvalue decomposition of estimated covariance matrices under the common effect null ($\eta = -0.33$), default initialization

| | $\lambda_1$ | $\lambda_2$ | $\lambda_3$ | $\lambda_4$ | Effective Rank |
|---|---|---|---|---|---|
| *(A) Genetic covariance* $\mathbf{C}_0$ | | | | | |
| Mean | 1.100 | 0.000 | 0.000 | 0.000 | 1.000 |
| Std | 0.484 | 0.000 | 0.000 | 0.000 | 0.000 |
| *(B) Error covariance* $\mathbf{C}_1$ | | | | | |
| Mean | 1.437 | 0.672 | 0.478 | 0.141 | 2.691 |
| Std | 0.386 | 0.104 | 0.050 | 0.119 | 0.353 |

$\lambda_i$: $i$-th eigenvalue (descending), $4 \times 4$ matrices, $n = 1{,}000$ reps; values below machine precision ($\sim 10^{-16}$) are reported as $0.000$.

**Table S9** Eigenvalue decomposition of estimated covariance matrices under the interaction effect null ($\eta = 1.0$), default initialization

| | $\lambda_1$ | $\lambda_2$ | $\lambda_3$ | $\lambda_4$ | Effective Rank |
|---|---|---|---|---|---|
| *(A) Genetic covariance* $\mathbf{C}_0$ | | | | | |
| Mean | 1.245 | 0.000 | 0.000 | 0.000 | 1.000 |
| Std | 0.648 | 0.000 | 0.000 | 0.000 | 0.000 |
| *(B) Error covariance* $\mathbf{C}_1$ | | | | | |
| Mean | 1.557 | 0.487 | 0.406 | 0.132 | 2.426 |
| Std | 0.580 | 0.047 | 0.040 | 0.101 | 0.455 |

$\lambda_i$: $i$-th eigenvalue (descending), $4 \times 4$ matrices, $n = 1{,}000$ reps; values below machine precision ($\sim 10^{-16}$) are reported as $0.000$.

**Table S10** Eigenvalue decomposition of estimated covariance matrices under the common effect null ($\eta = -0.33$), QR-based initialization

| | $\lambda_1$ | $\lambda_2$ | $\lambda_3$ | $\lambda_4$ | Effective Rank |
|---|---|---|---|---|---|
| *(A) Genetic covariance* $\mathbf{C}_0$ | | | | | |
| Mean | 1.589 | 0.524 | 0.252 | 0.080 | 2.088 |
| Std | 0.366 | 0.158 | 0.089 | 0.074 | 0.333 |
| *(B) Error covariance* $\mathbf{C}_1$ | | | | | |
| Mean | 0.775 | 0.408 | 0.229 | 0.066 | 2.585 |
| Std | 0.246 | 0.100 | 0.101 | 0.086 | 0.450 |

$\lambda_i$: $i$-th eigenvalue (descending), $4 \times 4$ matrices, $n = 1{,}000$ reps.

**Table S11** Eigenvalue decomposition of estimated covariance matrices under the interaction effect null ($\eta = 1.0$), QR-based initialization

| | $\lambda_1$ | $\lambda_2$ | $\lambda_3$ | $\lambda_4$ | Effective Rank |
|---|---|---|---|---|---|
| *(A) Genetic covariance* $\mathbf{C}_0$ | | | | | |
| Mean | 1.926 | 0.345 | 0.188 | 0.058 | 1.674 |
| Std | 0.506 | 0.097 | 0.068 | 0.057 | 0.268 |
| *(B) Error covariance* $\mathbf{C}_1$ | | | | | |
| Mean | 0.833 | 0.331 | 0.198 | 0.060 | 2.394 |
| Std | 0.361 | 0.071 | 0.080 | 0.074 | 0.428 |

$\lambda_i$: $i$-th eigenvalue (descending), $4 \times 4$ matrices, $n = 1{,}000$ reps.

#### Detailed Manhattan plots of joint GWAS

Per-chromosome Manhattan plots for *aINV* and *nINV* activity are provided in Figures 2 and 3, respectively, with SNPs colored by their relationship to loci reported by Fusari et al. (2017) and zoomed insets highlighting regions of high association density. Below we provide detailed per-locus results for each trait.

For *aINV*, four confirmed SNPs exceeded the Bonferroni threshold (LOD $\geq$ 6.63) under LRT10: *m7011* (LOD = 8.37) and *m7013* (LOD = 6.98) near *AT1G12240*, *m38695* (LOD = 8.18) near *AT1G62710* (Figure 2a), and *m141380* (LOD = 6.73) near *AT4G14368* (Figure 2d). Of the 14 additional SNPs (Supplementary Data S3, Additional Files), one (SNP 7009, LOD = 4.98) was detected under the common effect test (LRT10) within an existing QTL region near *AT1G12240*. The remaining 13 map to seven additional QTL regions: four on chromosome 1 and three on chromosome 5. Three were detected exclusively through the G$\times$E interaction test (LRT21): QTLs 55 and 79 (each one SNP, both LOD $\approx$ 4.85) and QTL 302 (two SNPs, both LOD = 4.81; Figure 2a,e). One region, QTL 71 on *AT1G51680* (one SNP, LOD = 5.35), was detected exclusively by the any effect test (LRT20; Figure 2a). The remaining three were detected by both the any effect and any vs. common effect tests: QTL 80 on *AT1G56240* (one SNP, LOD = 6.41 under LRT21), QTL 303 on *AT5G19300* (six SNPs, lead SNP 172599 with LOD = 6.45 under LRT21), and QTL 304 on *AT5G19300* (one SNP, LOD = 5.69 under LRT21), with lead SNPs in QTLs 80 and 303 approaching the Bonferroni threshold (Figure 2a,e). On chromosome 5, QTLs 302–304 clustered within a $\sim$3 kb region spanning *AT5G19290*–*AT5G19300* (Figure 2e).

For *nINV*, one confirmed SNP exceeded the Bonferroni threshold: *m196411* near *AT5G44560* (LOD = 7.11, LRT10; Figure 3e). *m16610* near *AT1G27720* (LOD = 6.61, LRT20; LOD = 6.23, LRT10; Figure 3a) approached but did not reach Bonferroni significance. The remaining two, *m115042* near *AT3G49430* (LOD = 4.46, LRT10; Figure 3c) and *m141380* near *AT4G14368* (LOD = 4.04, LRT10; Figure 3d), were recovered at moderate significance. Of the 23 additional SNPs, one (SNP 13405, LOD = 5.87) falls within the existing QTL region near *AT1G27720* and was detected under both LRT10 and LRT20. The remaining 22 map to 18 additional QTL regions (Supplementary Data S3, Additional Files). QTLs 24, 26–28 (five SNPs) span approximately 1.9 kb near *AT1G27720* and were detected through LRT20, with QTLs 24 and 26 also reaching significance under LRT10 (Figure 3a), while QTLs 42–48 and QTL 53 (11 SNPs) span approximately 13.3 kb near *AT1G35530*–*AT1G35537* and were detected predominantly through LRT21 (lead SNP 18887, LOD = 5.62; Figure 3a). The remaining six QTL regions are distributed individually across chromosomes 2, 3, and 5 (Figure 3b, c, e): QTL 121 near *AT2G23070* (SNP 54148, LOD = 4.85, LRT10), QTL 194 near *AT3G42180* (SNP 86116, LOD = 4.97, LRT20), QTL 217 near *AT3G45580* (SNP 90121, LOD = 5.39, LRT10; LOD = 5.25, LRT20), QTL 306 near *AT5G08060* (SNP 134082, LOD = 4.84, LRT20), QTL 327 near *AT5G43660* (SNP 158149, LOD = 5.15, LRT20; LOD = 5.03, LRT10), and QTL 340 near *AT5G51400* (SNP 164466, LOD = 4.82, LRT21). Overall, eight of the 18 additional QTL regions were detected exclusively through LRT21, four exclusively through LRT20, one exclusively through LRT10, and five through multiple tests. None reached Bonferroni significance.

Across both traits, 16 unique genes were tagged by the additional SNPs (LOD $\geq$ 4.8): 6 from aINV (AT1G34480, AT1G51680, AT1G55970, AT1G56240, AT5G19290, AT5G19300) and 8 from nINV (AT1G35530, AT1G35537, AT2G23070, AT3G42180, AT3G45580, AT5G08060, AT5G43660, AT5G51400) represent candidate genes not previously reported by Fusari et al., while 2 (AT1G12240 from aINV and AT1G27720 from nINV) fall within existing Fusari QTL regions (Supplementary Data S3, Additional Files). No additional genes were shared between both traits.

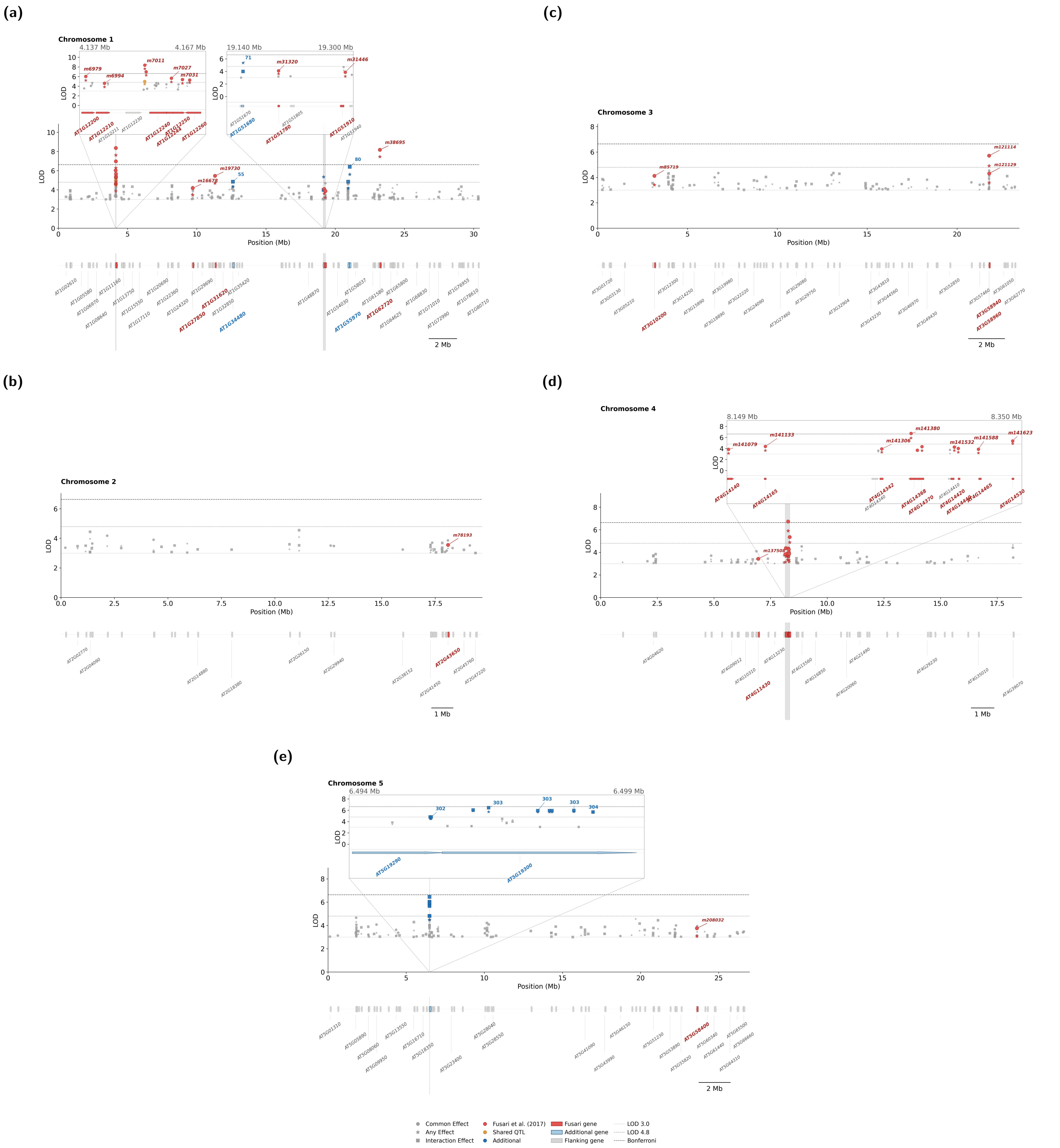


**Figure 2** Per-chromosome Manhattan plots for *aINV* activity across *A. thaliana* accessions. Each panel shows LOD scores $(-\log_{10}(p))$ for each SNP under three complementary tests: common effect (circles), any effect (stars), and any vs. common effect (squares) along chromosomal position (Mb). Highlighted SNPs either overlap with loci reported by Fusari et al. (2017) (red), represent additional SNPs sharing a QTL with a Fusari locus (orange), or represent additional associations above the significance threshold (blue). Dashed horizontal lines indicate LOD thresholds of 3.0, 4.8, and 6.63 (Bonferroni). The lower track in each panel shows gene models within association windows of SNPs exceeding LOD 3.0; labeled genes include those containing a significant SNP within their gene body, Fusari et al. (2017) candidate genes (red), genes in proximity to additional SNPs (blue), and flanking genes within the association window. To avoid crowding, gene labels are prioritized by the LOD score of the nearest associated SNP and suppressed if they fall within 400 kb of a higher-priority label. Zoomed insets highlight high-density association clusters. **(a)** Chromosome 1. **(b)** Chromosome 2. **(c)** Chromosome 3. **(d)** Chromosome 4. **(e)** Chromosome 5.

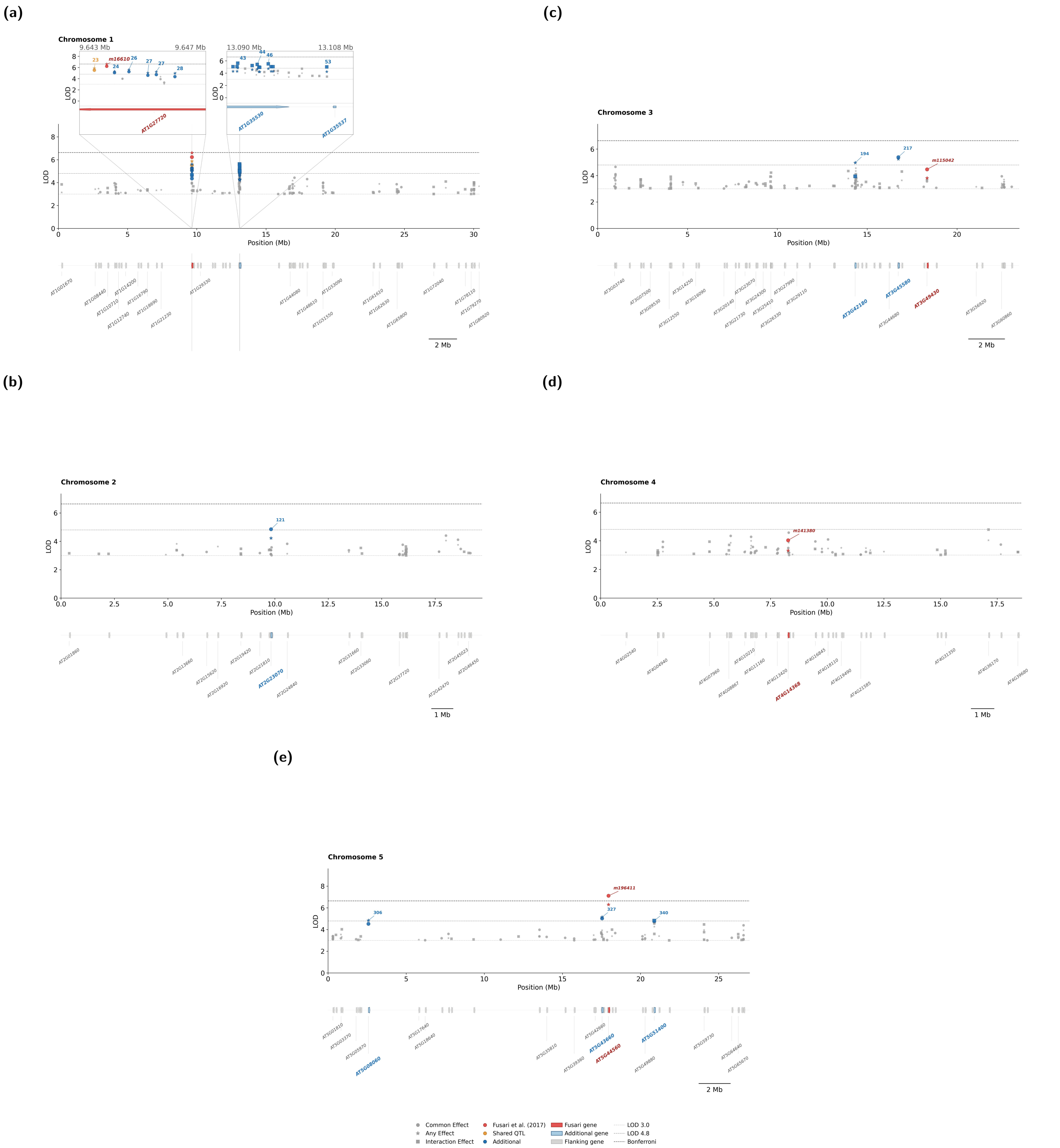


**Figure 3** Per-chromosome Manhattan plots for *nINV* activity across *A. thaliana* accessions. Each panel shows LOD scores ($-\log_{10}(p)$) for each SNP under three complementary tests: common effect (circles), any effect (stars), and any vs. common effect (squares) along chromosomal position (Mb). Highlighted SNPs either overlap with loci reported by Fusari et al. (2017) (red), represent additional SNPs sharing a QTL with a Fusari locus (orange), or represent additional associations above the significance threshold (blue). Dashed horizontal lines indicate LOD thresholds of 3.0, 4.8, and 6.63 (Bonferroni). The lower track in each panel shows gene models within association windows of SNPs exceeding LOD 3.0; labeled genes include those containing a significant SNP within their gene body, Fusari et al. (2017) candidate genes (red), genes in proximity to additional SNPs (blue), and flanking genes within the association window. To avoid crowding, gene labels are prioritized by the LOD score of the nearest associated SNP and suppressed if they fall within 400 kb of a higher-priority label. Zoomed insets highlight high-density association clusters. **(a)** Chromosome 1. **(b)** Chromosome 2. **(c)** Chromosome 3. **(d)** Chromosome 4. **(e)** Chromosome 5.

#### MapMan enrichment

MapMan bin enrichment analysis was performed separately for *aINV* and *nINV* using the nearest gene (within a ±10 kb window) for each SNP with LOD $\geq 3$. Over-representation was assessed by Fisher's exact test with Benjamini–Hochberg FDR correction applied independently at each hierarchy level (Supplementary Data S3 and S4). Results at levels 4 and 5 are summarized in Table S12 for *aINV*. For *nINV*, no bins reached nominal significance at level 4, and level 5 results are shown in Table S13.

**Table S12** MapMan bin enrichment analysis for acid invertase (aINV) activity (Levels 4 and 5). Fisher's exact test for over-representation among 412 genes with LOD > 3.0 versus 34,052 background genes. FDR correction: Benjamini–Hochberg per level. Only bins with nominal significance ($p < 0.05$) are shown; full results are available in Supplementary Data S4 (Additional Files).

| Bincode | Top-level category | Sig. genes | BG genes | Fold | Odds ratio | $p$-value | FDR |
|---|---|---|---|---|---|---|---|
| *Level 5* | | | | | | | |
| **2.2.1.3.3** | **major CHO metabolism** | **2** | **2** | **82.65** | $\infty$ | $\mathbf{1.46 \times 10^{-4}}$ | $\mathbf{1.61 \times 10^{-3}}$ |
| 29.5.11.4.3 | protein | 16 | 790 | 1.67 | 1.72 | $3.24 \times 10^{-2}$ | $1.31 \times 10^{-1}$ |
| 13.1.4.4.3 | amino acid metabolism | 1 | 3 | 27.55 | 40.92 | $3.59 \times 10^{-2}$ | $1.67 \times 10^{-1}$ |
| *Level 4* | | | | | | | |
| 2.2.1.3 | major CHO metabolism | 2 | 16 | 10.33 | 11.72 | $1.57 \times 10^{-2}$ | $1.53 \times 10^{-1}$ |
| 16.1.4.6 | secondary metabolism | 1 | 2 | 41.33 | 81.85 | $2.41 \times 10^{-2}$ | $1.53 \times 10^{-1}$ |
| 2.2.2.2 | major CHO metabolism | 1 | 2 | 41.33 | 81.85 | $2.41 \times 10^{-2}$ | $1.53 \times 10^{-1}$ |
| 16.1.1.1 | secondary metabolism | 1 | 3 | 27.55 | 40.92 | $3.59 \times 10^{-2}$ | $1.53 \times 10^{-1}$ |
| 17.6.1.12 | hormone metabolism | 1 | 3 | 27.55 | 40.92 | $3.59 \times 10^{-2}$ | $1.53 \times 10^{-1}$ |
| 29.4.1.51 | protein | 1 | 3 | 27.55 | 40.92 | $3.59 \times 10^{-2}$ | $1.53 \times 10^{-1}$ |
| 29.5.11.4 | protein | 23 | 1,287 | 1.48 | 1.51 | $4.18 \times 10^{-2}$ | $1.53 \times 10^{-1}$ |
| 9.2.1.2 | mitochondrial electron transport | 1 | 4 | 20.66 | 27.28 | $4.75 \times 10^{-2}$ | $1.53 \times 10^{-1}$ |

Bold = FDR $< 0.05$. Bin 2.2.1.3.3 corresponds to vacuolar invertases. Bin 2.2.1.3 is the parent category (sucrose degradation via invertases). FDR corrected independently per level.

**Table S13** MapMan bin enrichment analysis for neutral invertase (nINV) activity (Level 5). Fisher's exact test for over-representation among 280 genes with LOD > 3.0 versus 34,052 background genes. FDR correction: Benjamini–Hochberg. No bins reached FDR $< 0.05$. Only bins with nominal significance ($p < 0.05$) are shown; full results are available in Supplementary Data S5 (Additional Files).

| Bincode | Top-level category | Sig. genes | BG genes | Fold | Odds ratio | $p$-value | FDR |
|---|---|---|---|---|---|---|---|
| 29.5.11.4.2 | protein | 10 | 469 | 2.59 | 2.69 | $5.79 \times 10^{-3}$ | $5.79 \times 10^{-2}$ |
| 2.2.1.3.3 | major CHO metabolism | 1 | 2 | 60.81 | 121.04 | $1.64 \times 10^{-2}$ | $8.19 \times 10^{-2}$ |
| 13.1.1.1.1 | amino acid metabolism | 1 | 6 | 20.27 | 24.21 | $4.83 \times 10^{-2}$ | $9.99 \times 10^{-1}$ |
| 29.2.2.3.4 | protein | 1 | 6 | 20.27 | 24.21 | $4.83 \times 10^{-2}$ | $1.21 \times 10^{-1}$ |

Bin 2.2.1.3.3 corresponds to vacuolar invertases.
